# The rostral zona incerta: a subcortical integrative hub and potential DBS target for OCD

**DOI:** 10.1101/2022.07.08.499393

**Authors:** Suzanne N. Haber, Julia Lehman, Chiara Maffei, Anastasia Yendiki

## Abstract

**Background:** The zona incerta (ZI) is involved in mediating survival behaviors and is connected to a wide range of cortical and subcortical structures, including key basal ganglia nuclei. Based on these connections and their links to behavioral modulation, we propose the ZI is a connectional hub for in mediating between top-down and bottom-up control and a possible target for deep brain stimulation for obsessive compulsive disorder.

**Methods:** We analyzed the trajectory of cortical fibers to the ZI in nonhuman and human primates, based on tracer injections in monkeys and high-resolution diffusion MRI in humans. The organization of cortical and subcortical connections with the ZI were identified in the nonhuman primate studies.

**Results:** Monkey anatomic data and human dMRI data showed a similar trajectory of fibers/streamlines to the ZI. PFC/ACC terminals all converge within the rostral ZI (ZIr), with dorsal and lateral areas most prominent. Motor areas terminate caudally. Dense subcortical reciprocal connections included the thalamus, medial hypothalamus, substantia nigra/ventral tegmental area, reticular formation, and pedunculopontine nucleus and a dense nonreciprocal projection to the lateral habenula (LHb). Additional connections included amygdala, dorsal raphe nucleus, and periaqueductal grey.

**Conclusions:** Dense connections with dorsal and lateral PFC/ACC cognitive control areas and LHb and SN/VTA coupled with inputs from the amygdala, hypothalamus, and brainstem, suggests that the ZIr is a subcortical hub positioned to modulate between top-down and bottom-up control. A DBS electrode placed in the ZIr would involve both connections common to other DBS sites, but also would capture several critically distinctive connections.

## Introduction

The prefrontal/anterior cingulate cortex (PFC/ACC) is central to the network(s) that underlies behavioral flexibility [1–9]. These cortical areas and their connections to the basal ganglia (BG), thought to be dysfunctional in most psychiatric disorders, are the most common therapeutic targets for network modulation in several illnesses, including obsessive-compulsive disorder (OCD), mood disorders, and addiction. For example, the most common deep brain stimulation (DBS) targets for patients with treatment-resistant OCD are designed to capture descending PFC/ACC connections via the ventral anterior limb of the internal capsule (ALIC), the ventral striatum (VS), or anteromedial subthalamic nucleus (amSTN) [10–16]. However, this focus on the cortical-basal ganglia-thalamic system ignores other potential brain targets, specifically subcortical areas that also play important roles in modulating behavior in response to a rapidly changing environment. Given that DBS has been effective in only approximately 60% of patients [17], exploring the connectivity of targets outside the ‘light under the cortico-basal ganglia lamppost’, might, not only identify additional potential targets, but also increase our understanding of how circuits link across networks. Recent advances in imaging technology and of network analysis have identified cortical regions (i.e. ACC), referred to as hubs, with unusually high and diverse connections [18]. Hubs are thought to facilitate and integrate communication between different functional regions [19, 20, 21]. The emphasis has been on cortical hubs, yet, subcortical hubs are likely to be as critical for integration across functional domains. Here, we propose that the rostral ZI is subcortical hub positioned to modulate motivation and behavioral flexibility based on its PCC/ACC connections and links to basal ganglia structures, and brainstem region. As such, it may be potential DBS target.

The zona incerta (ZI), a small subcortical structure that lies dorsal to the STN and embedded within the fields of Forel, begins rostral to the mammillary bodies and extends caudal to the STN (Fig. 1) [22]. Rodent studies demonstrate its involvement in a large range of survival behaviors including fear and defense reactions, novelty, motivational drive, sensory integration, and motor control [23–36]. It is generally divided into a ventral/caudal (ZIv) and a dorsal/rostral part (ZId), bounded by rostral and caudal sectors [37–39]. The ZIv, has tight connections to motor and sensory cortical and subcortical regions, and the The ZId, has stronger links to limbic/affective areas, [29, 39–52]. Overall, the ZI likely plays a role in behavioral modulation [26]. The anterior medial sector is involved in behaviors suggestive for a role motivation, and reinforcement learning [27, 33, 34]. The nonhuman primate (NHP) ZI is also divided into four regions, the anterior pole/rostral region (ZIr), the ZId, ZIv and a caudal region (ZIc). Similar to rodents, the ZIv and ZIc are linked to sensory and motor systems [22, 53–55] and plays a role in novelty seeking [56]. However, little is known about PFC/ACC inputs to the ZI or how its connections intersect with BG circuits. Finally, the ZI is not part of the cortico-BG system [57, 58]. However, rodent studies show inputs from the SN/VTA and the LHb. In contrast, no mention of LHb-ZI connection has been reported in the monkey and little attention has been paid to a connection with the SN/VTA[59–61].

**Figure 1.**
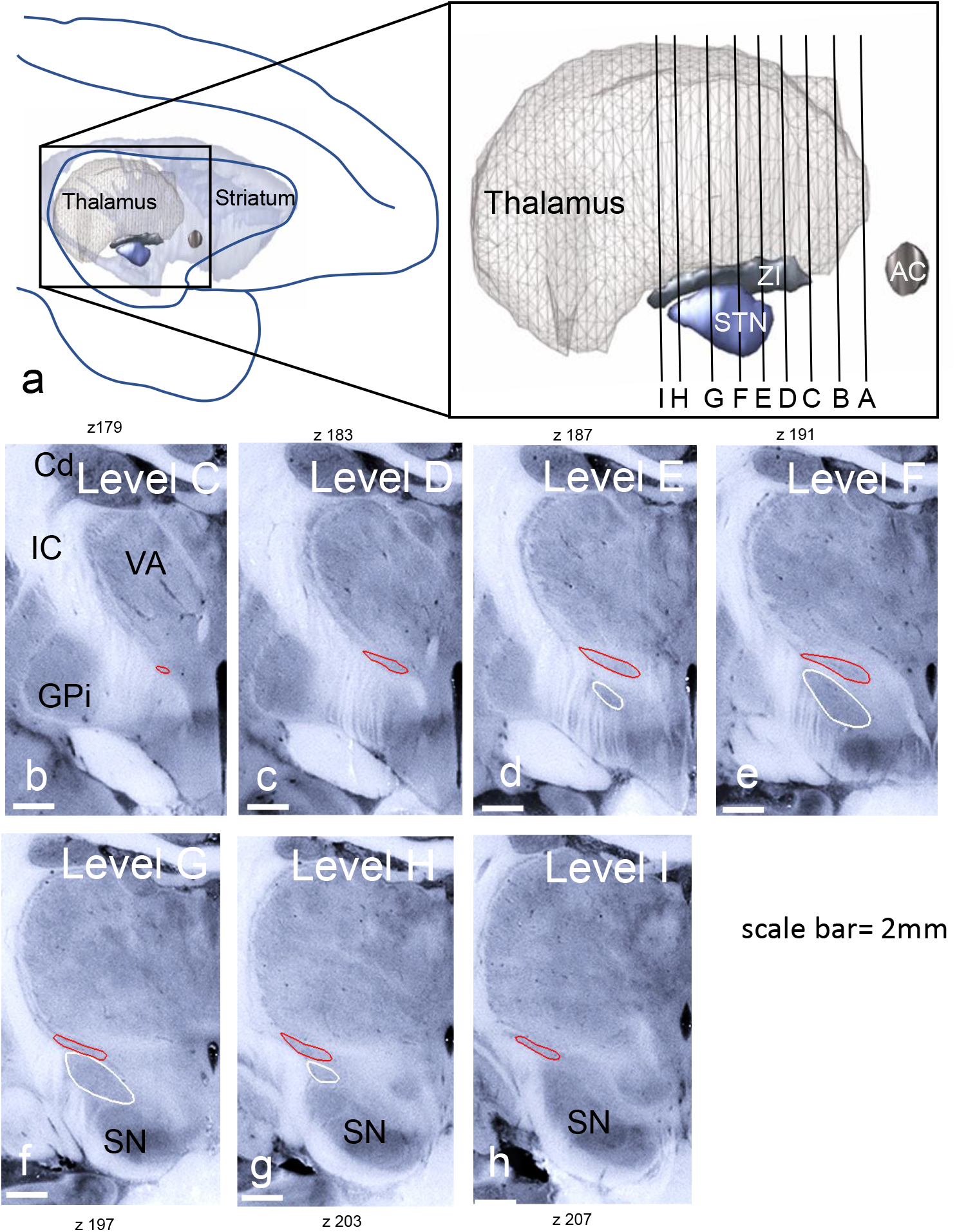
The position and extent of the ZI in the nonhuman primate. a. sagittal view illustrating the position of ZI and STN in context with the cortex, striatum and thalamus. Vertical lines shown in the blow-up indicate the coronal levels shown in b-h. b-h. Blockface coronal sections, illustrating the rostral-caudal position of the ZI (red) and STN (white). Coronal levels are indicated in the upper left for each section. ZIr=levels C-E, ZId & ZIv=levels E-H, ZIc=levels H-I.

DBS for OCD primarily targets OFC/ventral ACC-basal ganglia-thalamic connections through the ventral ALIC, VS, and amSTN [13, 16, 62, 63]. At the typical parameters used for stimulation, myelinated fibers are preferentially stimulated over nonmyelinated fibers and terminals [64]. Thus, one aim of this study was to outline the trajectory of fibers from each frontal cortical area through the internal capsule (IC), with the specific goal of identifying their branching points to the thalamus, ZI, STN, and brainstem. Using diffusion MRI tractography at sub-mm resolution, we then demonstrated these branch points in the human. Our second aim was to identify the organization of PFC/ACC terminals in the ZI compared to motor control regions. The third aim was to delineate other connections to the region of the ZI that receives PFC/ACC inputs. The results show that the PFC/ACC projects primarily to the ZIr, with the strongest projections from cognitive control areas. They also demonstrate strong ZIr connections with two key BG areas, the LHb and SN/VTA. Additional connections to the ZIr include the amygdala, hypothalamus, PAG, DRN, PNN, and superior colliculus. PFC/ACC projections, coupled with links to the BG system and non-BG subcortical inputs indicate that the ZIr serves as a subcortical hub in which cognitive, top-down control can be exerted on a subcortical network primed for quick, survival-based responses to a changing environment. These findings are discussed in the context of a potential DBS target for OCD.

## Methods

### Anatomical tracing experiments

We used a combination of our extensive library of previous experiments, in which tracer injections were placed in different frontal cortical and subcortical areas, and 4 new experiments in which we specifically targeted the ZIr. All animals were adult male macaque monkeys (*Macaca nemestrina, Macaca fascicularis, and Macaca mulatta*). Three sets of injections sites were analyzed: 1. Cortical injections to determine the trajectory of fibers as they leave cortex, enter and exit the IC and the terminal fields in the ZI; 2. ZIr injections to determine its afferent and efferent connections; 3. Injections into the LHb to its ZIr connection. Anterograde or bidirectional tracers were injected into the following cortical areas, grouped based on six regions: ventromedial PFC (vmPFC)-areas 25, 14, 10, and subgenual ACC), rostral ACC (rACC)-areas 32 and pregenual 24), dorsal (dACC)-(area 24, dorsomedial PFC (dmPFC) medial area 9 and dorsal area 10, dorsolateral PFC (dlPFC)-areas 46 and lateral 9, ventrolateral PFC (vlPFC-areas 44, 45, and 12/47), and orbitofrontal cortex (OFC)-areas 11 and 13. Anterograde or bidirectional tracers were also placed in the ZIr. Finally, to evaluate specifically the connection of the ZIr, we analyzed three LHb injection sites. All experiments were carried out in accordance with the Institute of Laboratory Animal Resources Guide for the Care and Use of Laboratory Animals and approved by the University Committee on Animal Resources.

Details of the surgical and histological procedures have been described previously [65–67]. Briefly, monkeys received an injection of one or more of the following anterograde/bidirectional tracers: Lucifer yellow, fluororuby, or fluorescein conjugated to dextran amine (LY, FR, or FS; 40–50 nl, 10% in 0.1M phosphate buffer (PB), pH 7.4; Invitrogen) or tritiated amino acids (100 nl, 1:1 solution of [^3^H] leucine and [^3^H]-proline in dH2O, 200 mCi/ml, NEN). Tracers were pressure-injected over 10 min using a 0.5 μl Hamilton syringe. Following each injection, the syringe remained *in situ* for 20–30 min. Twelve to 14 days after surgery, animals were initially anaesthetized with Ketamine (10 mg ⁄ kg, intramuscularly) and then deeply anesthetized and perfused with saline, followed by a 4% paraformaldehyde/1.5% sucrose solution in 0.1 M PB, pH 7.4. Brains were postfixed overnight and cryoprotected in increasing gradients of sucrose (10%, 20%, and 30%). Serial sections of 50 um were cut on a freezing microtome and immunocytochemistry was performed on free-floating sections (1 in 8 for each tracer) to visualize LY, FR, and FS tracers. Sections were mounted onto gel-coated slides, dehydrated, defatted in xylenes, and coverslipped with Permount. To visualize amino acid staining, sections were mounted on chrome-alum gelatin-subbed slides for autoradiography. Sections were defatted in xylene for 1 h, and then dipped in Kodak NTB 2 photographic emulsion. Exposure time of the autoradiograms ranged from 6 to 9 weeks. The sections were then developed in Kodak D for 2.5 min, fixed, washed, and counterstained with cresyl violet.

#### Data Analysis

65 frontal cortical injection sites (out of 156) were selected for the initial analysis of cortico-ZI projections based on the following criteria: 1. localized injection site without contamination into adjacent cortical regions or into WM; 2. overall outstanding or excellent transport; and 3. low background. From these cases 28 were analyzed in detail based on outstanding transport to the ZI coupled with a systematic sampling of cortical areas (Fig. 2). Under darkfield microscopy, outlines of the brain sections and the fibers traveling in bundles were charted with a 4.0, or 6.4x objective respectively, using Neurolucida software (MBF Bioscience). Orientation was indicated for each bundle by charting individual fibers (at 10X) within each outline (see Fig. 3, c-i). Axons from each injection sites were charted as they left the tracer injection site and followed as they entered the ALIC, passed through, and the point at which they exited the IC to the thalamus, ZI and STN. The 2D outlines were combined across slices to create 3D renderings of the pathways using IMOD software (Boulder Laboratory for 3-Dimensional Electron Microscopy of Cells and the Regents of the University of Colorado) [68]. This was done for each case separately and then registered to a reference model to compare the relationship between fiber positions from the different cortical regions. For details of this method see [65]. In addition to the cortical tracer injections, four new injections were placed in the ZIr, three bidirectional tracers and one tritiated amino acid. Processing and analysis were the same as described above. However, in addition to terminal fields, labeled cells from the retrograde component of the bidirectional tracers were also charted. Emphasis was placed on the distribution of labeled cortical cells to verify the anterograde results from the cortical experiments. Finally, we used 4 injections placed in the LHb nucleus to validate the new finding of a ZIr connection to this nucleus.

**Figure 2.**
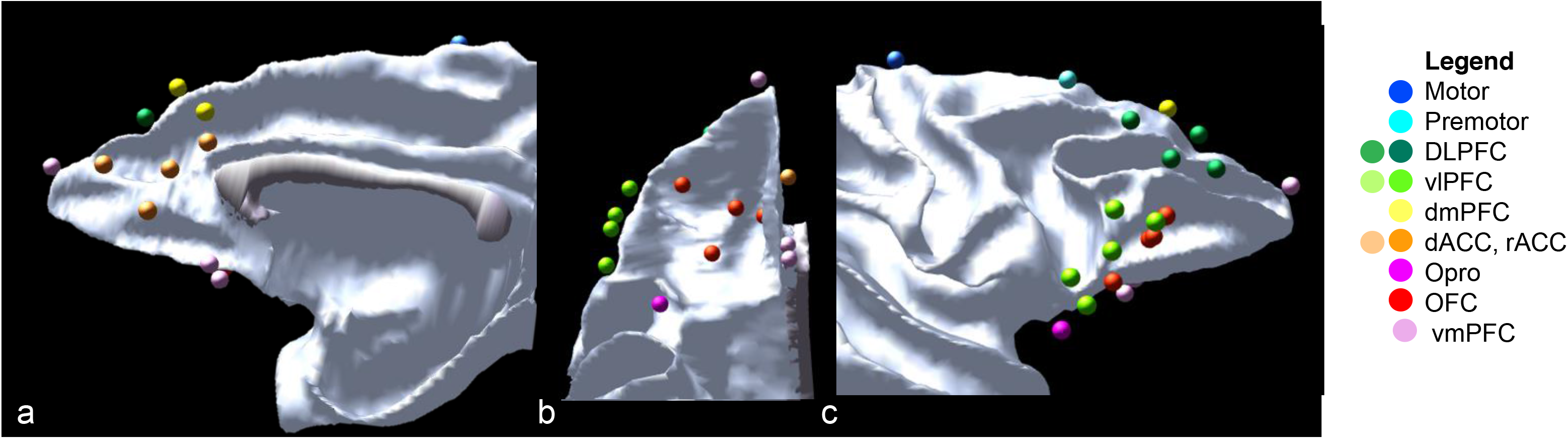
Location of cortical injection sites. Pink=vmPFC (areas 25 & 14), Red=OFC (areas 11 & 13), fuchsia=Opro, orange=rACC & dACC (areas 32 &24), yellow=dorsomedial PFC (area 9), light green=vlPFC (areas 45&47), dark green=dlPFC (areas 9 & 46), light blue=premotor (area 6), dark blue=motor (M1).

**Figure 3.**
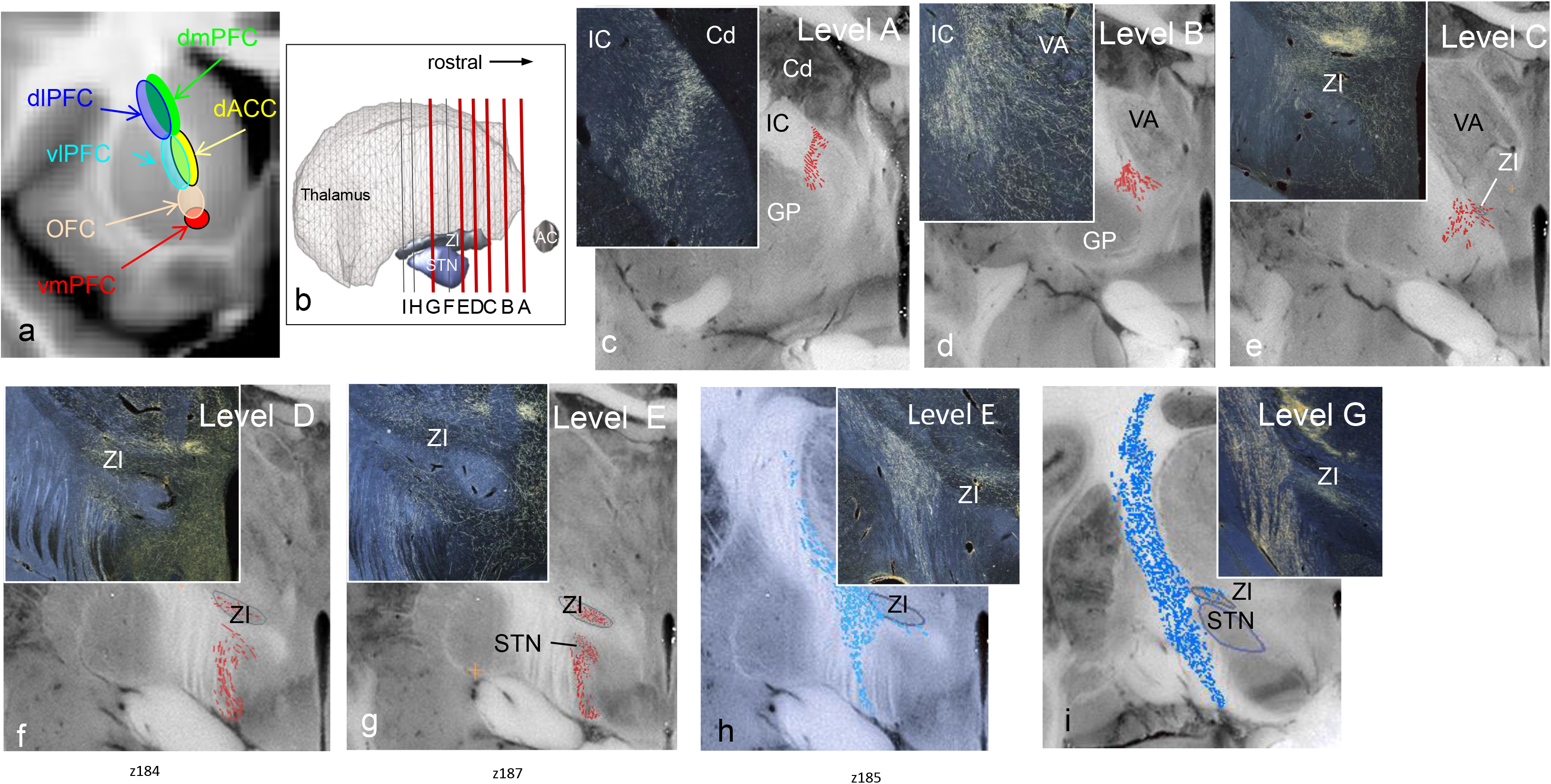
Frontal fibers as the travel through and exit the internal capsule to the ZI, STN, and brainstem. a. Schematic illustrating the dorsoventral, mediolateral topology of PCC/ACC fibers as they pass through the ALIC. b. Sagittal view of the thalamus, STN, and ZI. Red vertical lines =coronal levels shown in c-i. c-i. Coronal sections through different levels indicated in b, illustrating, the orientation of IC fibers as they exit to the thalamus, ZI, and STN. Each panel shows the orientation of fibers on blockface coronal level, red=PFC fibers, blue=motor fibers. Insets=photomicrograph examples for each level. c-g ACC fibers, h premotor, and i. M1 fibers.

### Diffusion MRI

*Acquisitio*n. Diffusion MRI (dMRI) data were acquired on the MGH-USC 3T Connectom Scanner at 0.76 mm isotropic resolution using an SNR-efficient simultaneous multi-slab imaging technique (gSlider-SMS) [69] [70]across 9 2-hour sessions (max gradient amplitude: 180 mT/m; slew rate: 125 T/m/s, gSlider factor: 5, MB factor: 2, R: 3, TR/TE: 3500/75 ms, matrix: 290x288, PE: AP). 2808 dMRI volumes were acquired (144 b=0, 420 b=1000, 840 b=2500 s/mm^2^, along with their paired reversed PE volumes). T1-weighted images were also acquired with the Multi-echo Magnetization-Prepared Rapid Acquisition Gradient Echo (MEMPRAGE) [71] at 1 mm isotropic resolution (TR=2350 ms; TI=1100 ms; number of echoes=4; TEs=1.61, 3.47, 5.34, 7.19 ms; flip angle=7; FOV=256x256x208 mm^3^; bandwidth=650Hz/Pixel; GRAPPA factor =2; total acquisition time=6 min).

#### Processing

We used pre-processed, publicly available data [72]. We fit fiber orientation distribution functions (fODFs) to the dMRI data using multi-shell multi-tissue constrained spherical deconvolution (MSMT-CSD [73]) n MRtrix3 [74]. We performed probabilistic tractography with whole-brain dynamic seeding to obtain 20M streamlines. The following tractography parameters were used: step-size = 0.38 mm, maximum angle threshold = 45°, maximum length = 150 mm. Cortical parcellations and subcortical segmentations were obtained from the T1 data using FreeSurfer[75–77] [78]. Segmentations of the thalamic nuclei were also obtained [79]. To maximize correspondence between human and NHP prefrontal areas, we used a publicly available parcellation scheme which translated the anatomical definitions of cytoarchitectonic regions from Petrides et al 2012[80] to the fsaverage cortical surface [81]. We mapped that parcellation from the fsaverage surface to the individual surface using the inverse of the FreeSurfer spherical morph. We then used a boundary-based, affine registration method [81] to align the T1 to the dMRI and map the parcellations and segmentations volumes to dMRI space.

#### Virtual dissections

Tracts were annotated manually by author C.M. in Trackvis (v.0.6.1; http://www.trackvis.org). To select only streamlines belonging to the internal capsule (IC), the IC was manually labeled by author C.M. on two axial and two coronal slices to include both its frontal and motor projections and used as inclusion region of interest (ROI). To isolate streamlines connecting the ZI and STN, these regions of interest (ROIs) were manually labeled in diffusion space by author S.H. on the CSD-based maps (volume correspondent to the 0-th spherical harmonic order) and used as inclusion ROIS. ZI connections were defined as streamlines passing through the ZI and the IC. STN connection were defined as streamlines passing through the STN and the IC. Connections to the thalamus were also isolated, using the FreeSurfer segmentation label of the entire thalamus. For all tracts, a sagittal exclusion ROI was placed on the midline to exclude streamlines erroneously crossing to the opposite hemisphere. No other main exclusion ROIs were used. A maximum length threshold of 100 mm was applied to the ZI, STN tracts to eliminate spurious streamlines. To further subdivide the streamlines based on their cortical projections, we used the labels from the Petrides parcellation to define the following cortical inclusion ROIs: motor (region 4); premotor (region 6); dorso-medial pre-frontal cortex (dmPFC; region 9M); dorso-lateral PFC (dlPFC; regions 946v, 946d, 46, 9L); ventro-lateral PFC (vlPFC; regions 47O, 47L, 45A, 45B, 44); (dACC; regions 24, 32); orbito-frontal cortex (OFC; regions 14O, 14M); Opro (region 13

## Results

### Frontal projections to the ZI

#### PFC/ACC fiber trajectories through the IC to the ZI, STN, and brainstem

PFC/ACC axons leave the grey matter and form a tight ‘stalk’ as they travel through the corona radiata to enter the ALIC [82, 83, 84]. Fibers from each cortical region remain clustered together and organized topologically as they pass ventro-caudally through the ALIC (Fig. 3a) [82]. However, caudal to the anterior commissure, just anterior to the emergence of the thalamic reticular nucleus, PFC/ACC fibers begin to spread across the mediolateral aspect of the ALIC, and begin to separate into two groups of axons (Fig. 3c): one orienting medially to enter the thalamus (via the anterior thalamic peduncle) and one that continues to travel ventrally within the capsule to the ZI, STN, hypothalamus, and brainstem. PFC/ACC axons traveling to the ZI split off from the IC as the ventral anterior thalamic nucleus forms and both the external and internal segments of the globus pallidus are evident (Fig. 3d). Fibers from the PFC/ACC enter the ZI as the ZIr begins to form (Fig. 3e). Axons traveling to the STN and brainstem remain as a bundle within the IC. At the emergence of the STN, PFC/ACC bundles split again, with a branch of fibers exiting the IC to rostral STN (Fig. 3f). Here some fibers travel caudo-medially, circumventing the STN to terminate in the SN/VTA, while others continue in the IC to lower brainstem regions (Fig. 3g). In contrast, fibers from premotor areas travel in the IC more caudally and enter the ZI at the level of the STN, but not in the ZIr (Fig. 3h). Fibers from motor cortex pass through the IC, and branch to thalamus, ZI, STN approximately at the same coronal level (Fig. 3i). Figure 4 compares the fiber trajectories from different frontal regions as they pass through the IC and branch to the ZI, STN, and SN/VTA. Taken together, PFC/ACC fibers enter the thalamus, ZI, and STN at approximately the same rostral level as the ZI forms (Fig. 4d-e). At this level, premotor fibers are positioned dorsally within the IC and motor fibers have not yet entered the IC (Fig. 4c-f).

**Figure 4.**
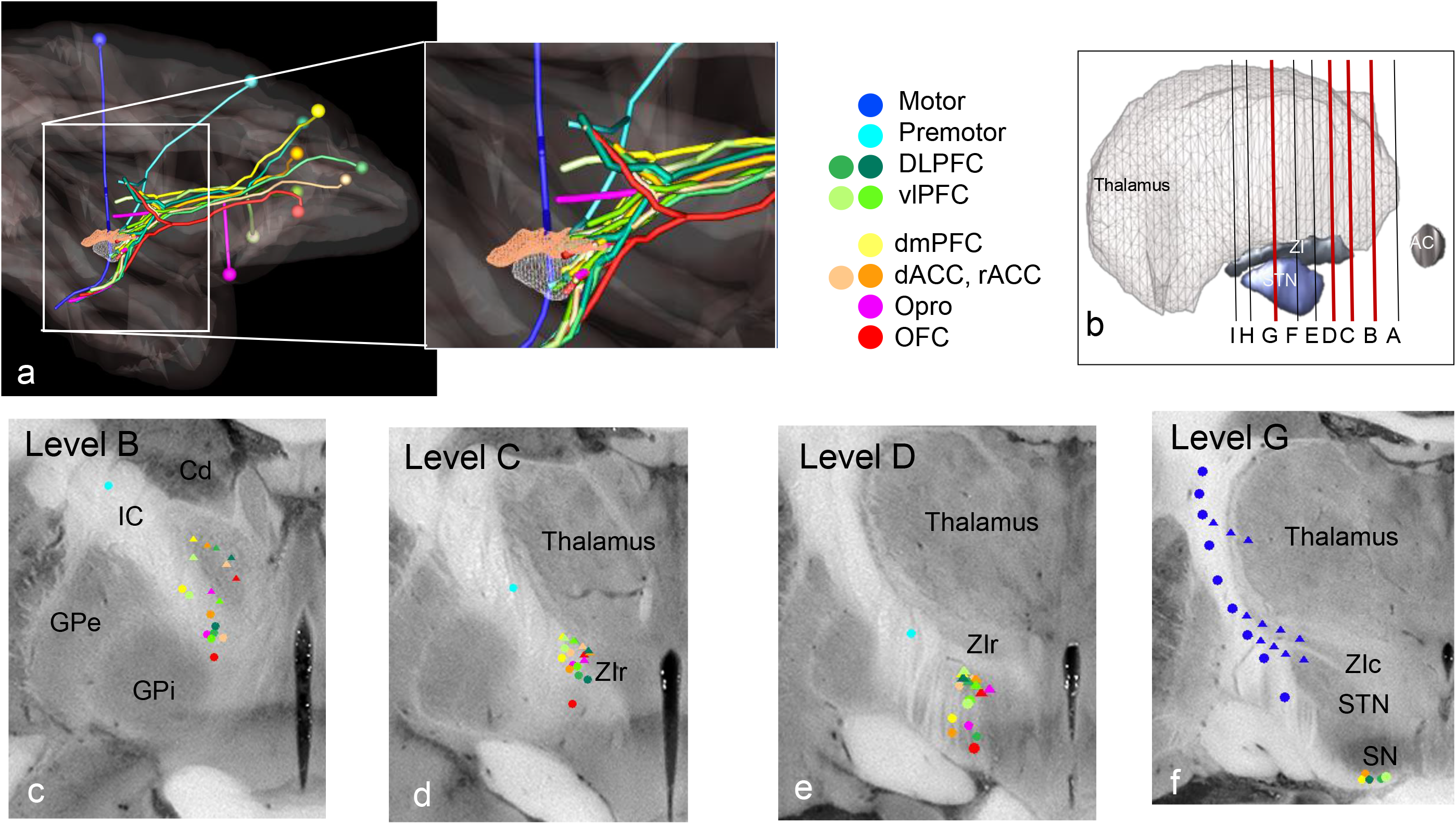
Schematic illustrating the trajectory of frontal fibers as enter and the course through and exit the IC to the thalamus, ZI, and STN. a. sagittal view illustrating the injection sites (balls), and fiber trajectories. Inset, illustrating the branching points for each set of cortical fibers as they first exit at the thalamus, then ZI and STN and continue to the brainstem. Red=OFC (areas 11 & 13), fuchsia=Opro, orange=rACC & dACC (areas 32 &24), yellow=dorsomedial PFC (area 9), light green=vlPFC (areas 45&47), dark green=dlPFC (areas 9 & 46), light blue=premotor (area 6), dark blue=motor (M1). b. schematic illustrating the levels for c-f. c-f. coronal blockface sections at levels B-D and G illustrating the position of frontal fibers within the IC and as they exit to the thalamus and ZI. Circles=passing through the IC, triangles=fibers exiting the IC to the thalamus, ZI, and STN.

Using diffusion MRI tractography in the human brain, streamlines could be correctly identified as they exit the IC to enter the ZI, STN, and SN (Fig. 5). Consistent with the tracing, streamlines begin to change orientation within the IC, rostrally at the level of the emerging thalamus, with those connecting to the dorsal thalamus orienting medially. Fig. 5b illustrates the change in orientation at approximately the same level as noted in the tracing experiments (see Fig. 3c). Caudally, streamlines exit the IC to enter the ZI as it emerges (Fig. 5c). This is approximately at a level similar to where the PFC/ACC fibers in tracer experiments exit the IC to the ZI (see Fig. 3d). Caudal to the ZI, streamlines exit the IC to enter the STN, while others continue inferiorly (Fig. 5d, e). In agreement with tracer data, streamlines originating from PFC regions enter the ZI at its most rostral aspect, while streamlines originating in pre-motor and motor regions enter the ZI posteriorly (Fig. 6). As such, as with the tracing data, there is a clear rostro-caudal organization of frontal connections to the ZI. PFC/ACC streamlines enter the ZI rostrally at the level in which motor areas are entering the thalamus (Fig. 6e). Caudally, as streamlines from motor regions enter the ZI, those from the PFC/ACC travel to the brainstem (Fig. 6f, g).

**Figure 5.**
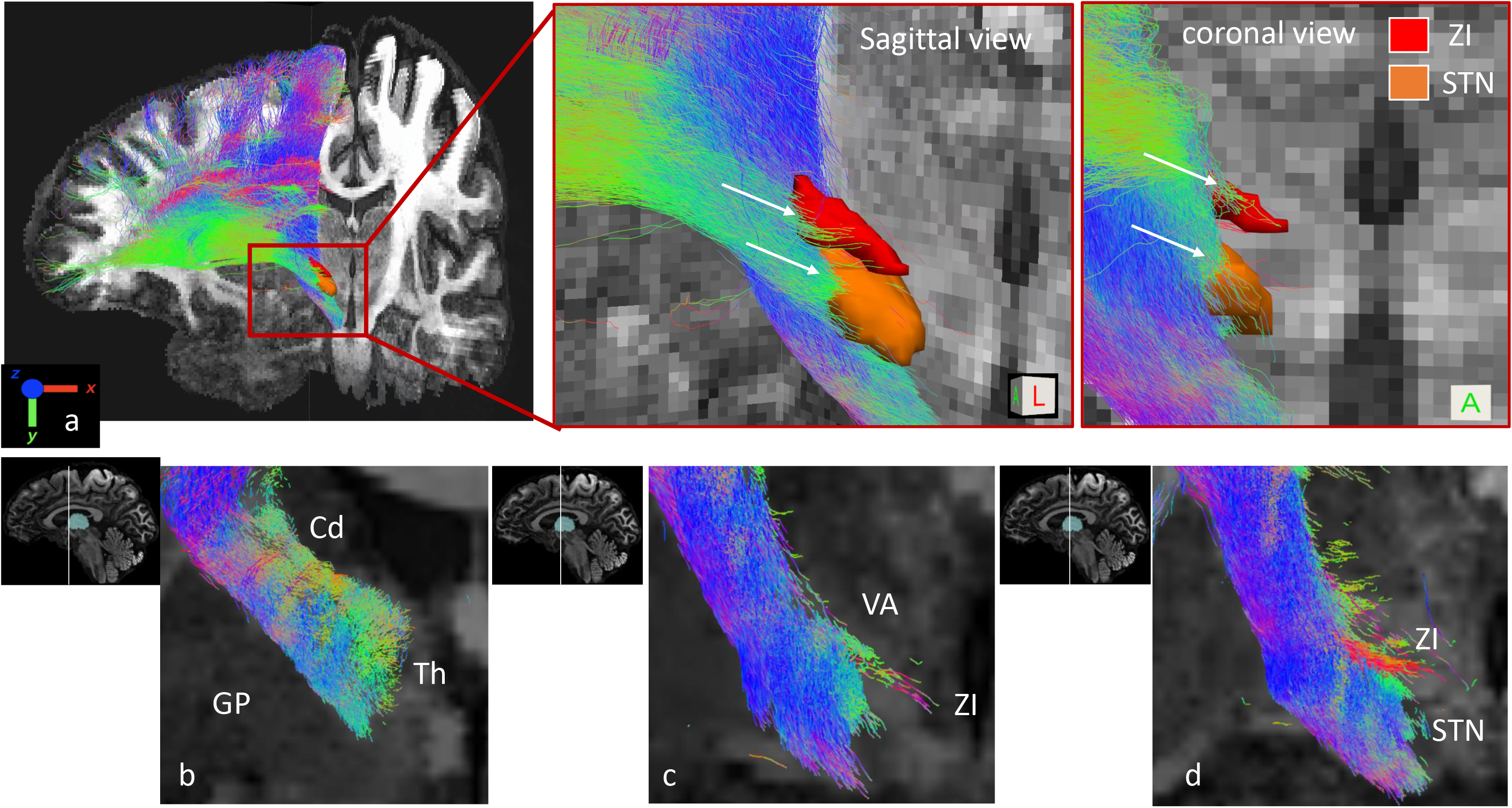
Frontal cortical (right hemisphere) streamlines as they course through and exit the IC to the ZI and STN. a. Fronto-lateral view of the streamlines coursing through the IC and entering the ZI (red) and STN (orange) (left panel). The red box=magnification area shown in the central (lateral view) and right panels (coronal view). The white arrows=streamlines leaving the IC to enter the ZI and STN. b-d. Coronal views showing the streamlines three different coronal levels to correspondent to the NHP anatomic levels shown in figure 3. Sagittal views next to each panel shows the location of the coronal slide with respect to the thalamus (shown in light blue). b. Streamlines change orientation as one branch exits the IC to the thalamus (approximately Level A in Fig. 3). c. Streamlines exit to the ZI (approximately Level B in Fig. 3). d. Streamlines continue into the ZI, other exit the IC to the STN (approximately Level C in Fig. 3). Color=direction; red=left-right, blue=inferior-superior, green=antero-posterior. Cd=caudate nucleus, GP=globus pallidus, STN=sub-thalamic nucleus, Th=thalamus, VA=ventral anterior nucleus of the thalamus, ZI= zona incerta.

**Figure 6.**
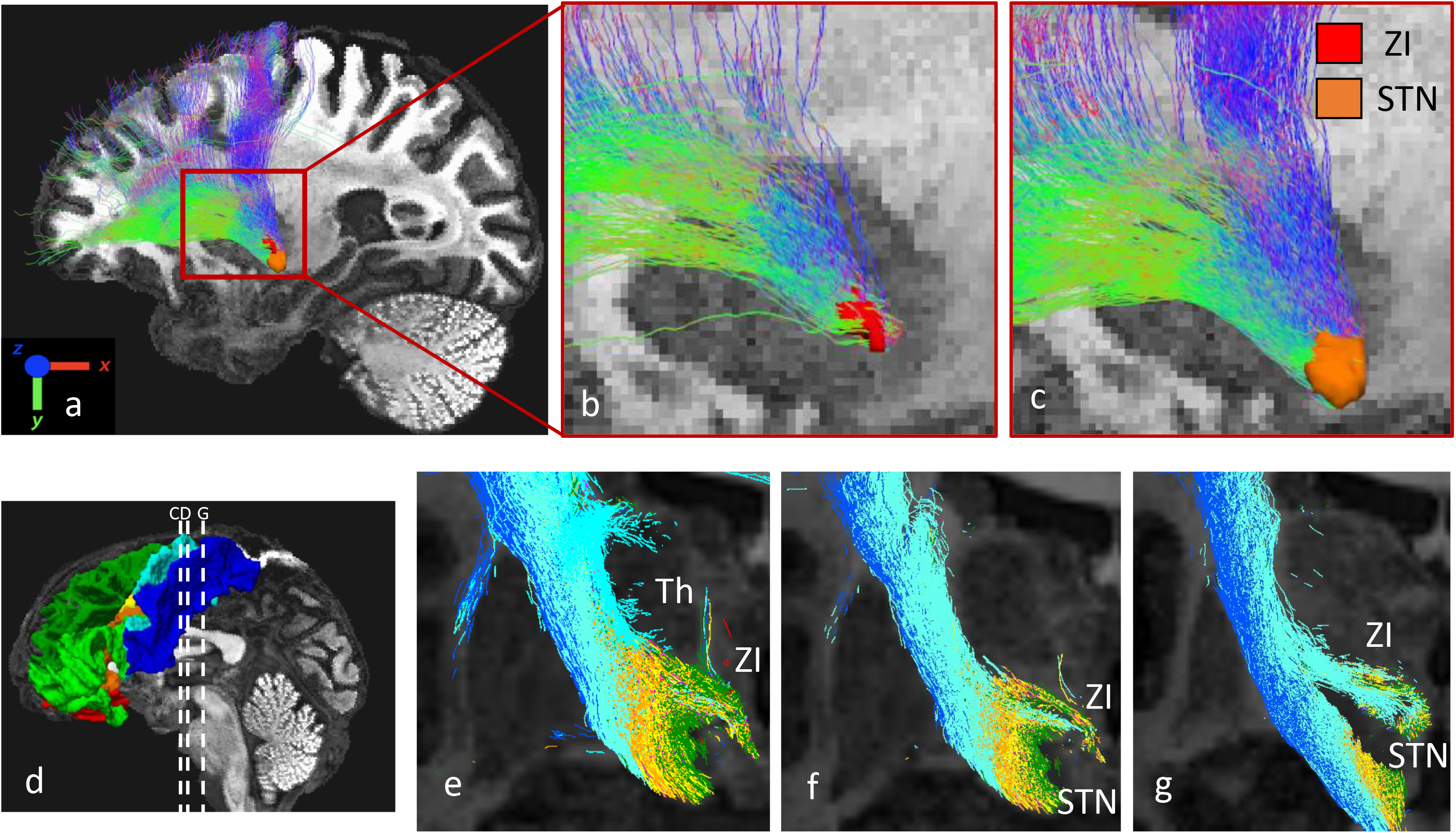
a. Lateral view of the streamlines entering the ZI (red) and STN (orange). Streamlines coursing inferior to enter the brainstem were excluded. b-c. magnification of area indicated in a. b. only streamlines entering the ZI. c. only the streamlines entering the STN. White curves indicate the separability of rostral and caudal projections. Streamlines color-coded as in Fig. 5). d-g. Topology of front-cortical fibers entering the ZI and STN color-codes and based on cortical areas. d. sagittal view of cortex, indicating coronal levels shown in d-g. levels correspond approximately to the levels C, D, G in Fig. 3. e-g. Streamlines are color-coded based on their cortical projections (see Fig. 4), blue=motor, light-blue=premotor; dark green=dlPFC; light green=vlPFC; yellow=dmPFC; orange=dACC; fuchsia=Opro; red=OFC. Cd: caudate nucleus; GP: globus pallidus; STN: sub-thalamic nucleus; Th: thalamus; VA: ventral anterior nucleus of the thalamus; ZI: zona incerta.

#### Frontal cortical terminals within the ZI

As PFC/ACC fibers exit the IC and enter the ZIr, they immediately begin to synapse as the nucleus forms (Fig. 7a-d). As such, terminals from all PFC/ACC converge at approximately the same level in the ZIr. However, many axons also continue to travel medially through the ZI giving off boutons en passage (Fig. 7e). PFC/ACC terminal fields are, therefore, also found throughout the mediolateral ZIr. The axons and terminals continue into the rostral part of the central ZI (Fig. 7f-h). The density of these projections varied based on the cortical region of origin. Those from area vmPFC were the sparsest (Fig. 7b, g). In contrast, axons originating from the rACC, dACC, dmPFC, dlPFC, vlPFC, and premotor control areas have relatively dense terminal fields (Figs. 7c-g & Fig. 8a-c). Fibers from premotor and motor areas enter and terminate in the central ZI (a combination of both ZId and ZIv) and ZIc (Fig. 8b, c). However, those from primary motor areas, M1, were remarkably sparse. Axons from motor areas also travel medially through the ZI and continue to the red nucleus (Fig. 8). Following bidirectional tracer injections in the rZI, retrogradely labeled cells are located primarily in layer V throughout much of frontal cortex. Consistent with the anterograde experiments, the labeled cells are located primarily in PFC/ACC and concentrated in the dlPFC, vlPFC, and the rostral and dorsal ACC (Fig. 9a-d), with few labeled cells in the ventromedial PFC regions. Moreover, the density of labeled cells decreased in caudal sections, as motor areas emerge (Fig. 9e). In summary, the ZI received dense inputs from cognitive and premotor areas, including ACC regions associated with both cognition and action planning, but relatively few inputs from primary motor areas and visceral limbic regions (subgenual ACC and medial OFC). There was a general rostro-caudal gradient of fronto-incerta projections: PFC/ACC projections were concentrated in the ZIr and projections from motor control areas in the ZIc. However, in the central ZI terminals from both PFC/ACC and premotor regions overlap.

**Figure 7.**
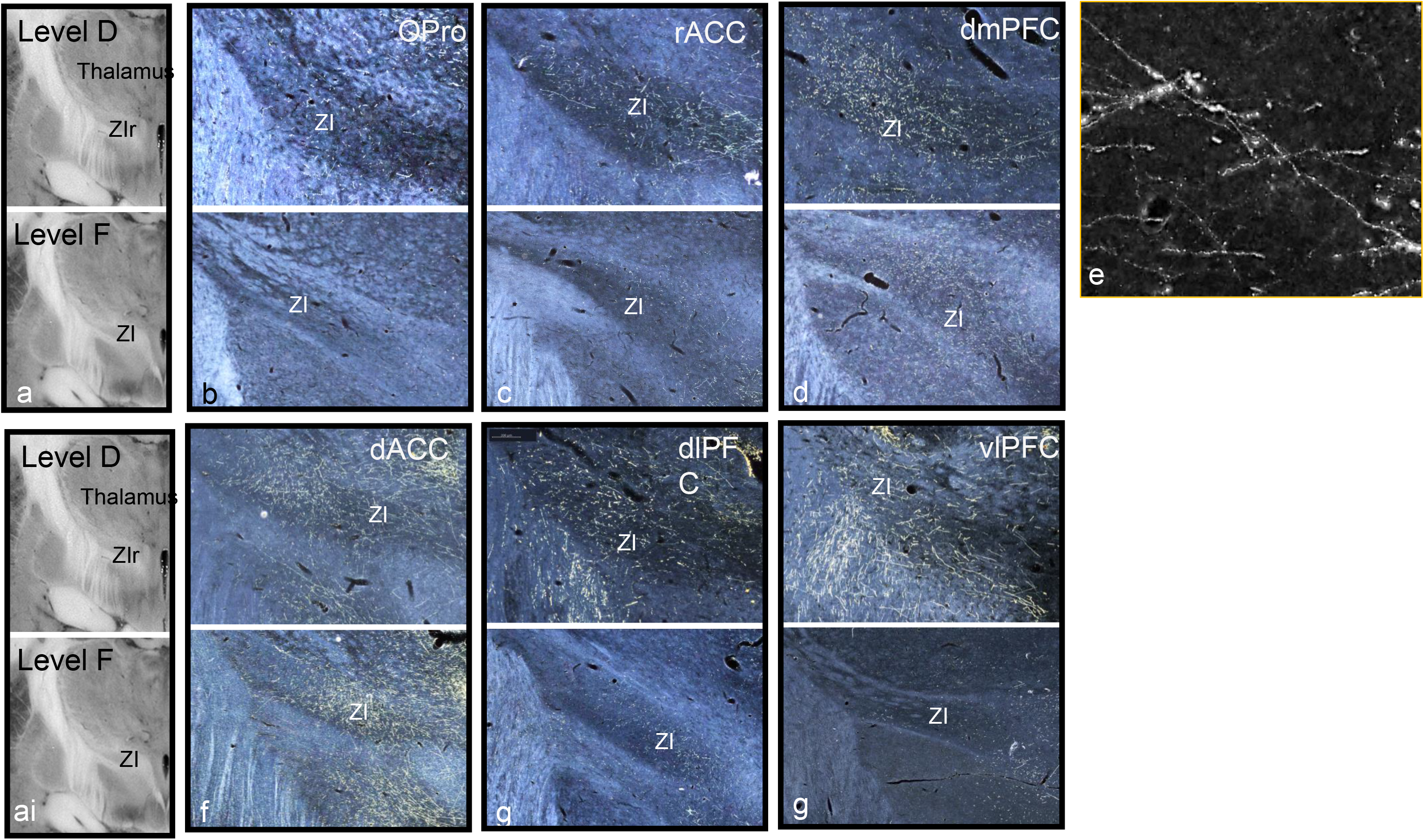
Photographs of PFC/ACC terminals in the ZI. a. & ai. blockface image of levels D & E illustrated in each row. b. terminals and fibers from OFC. c. terminals and fibers from rostral ACC. d. terminals and fibers from dorsomedial PFC. E. examples boutons en passage. f. terminals and fibers from dorsal ACC. g. terminals and fibers from dorsolateral PFC. h. terminals from ventrolateral PFC.

**Figure 8.**
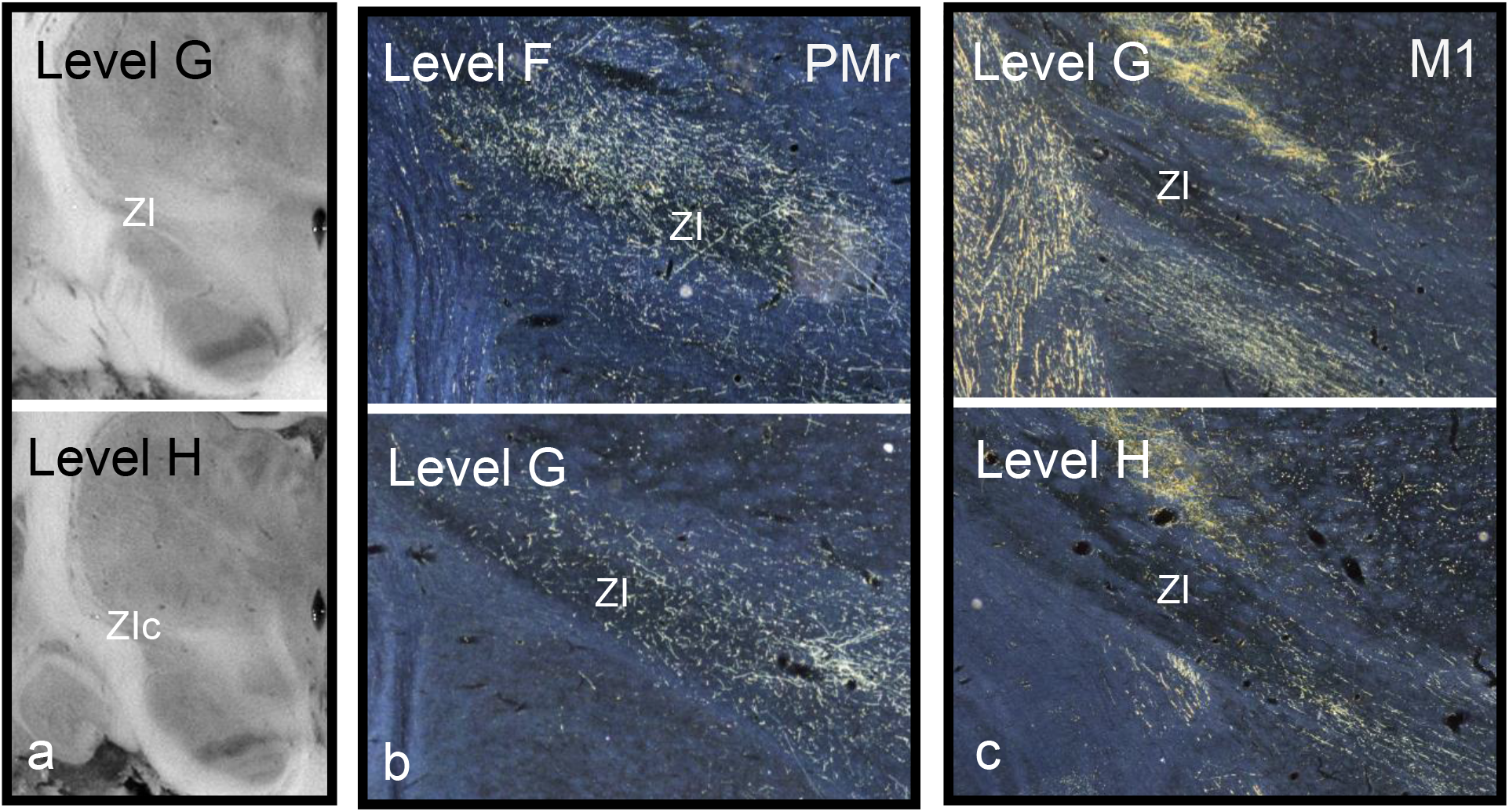
Photographs of premotor and motor terminals in the ZI. a. l blockface image of levels G & H illustrated in each row. b. terminals and fibers from premotor cortex. c. terminals and fibers from motor cortex.

**Figure 9.**
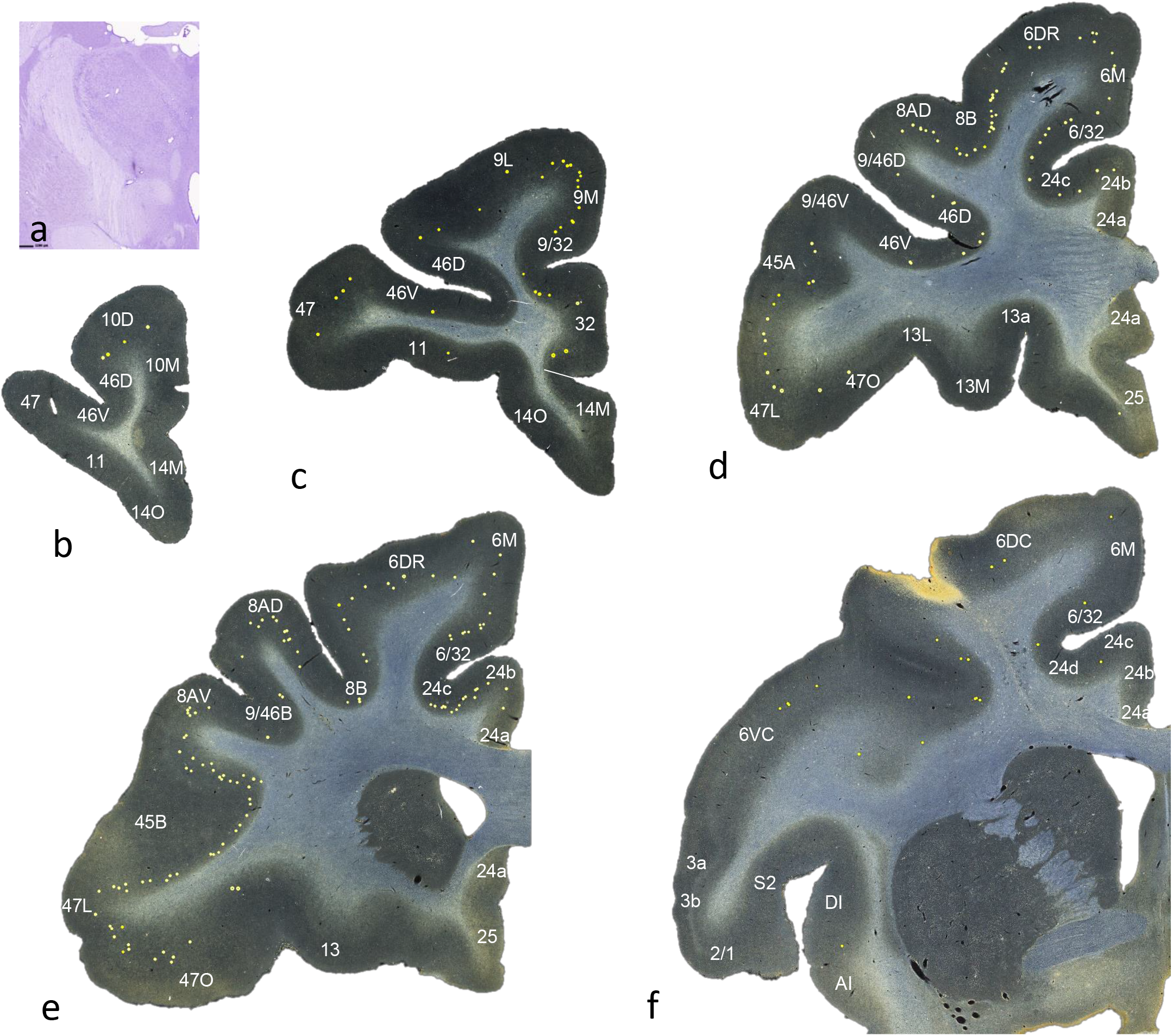
Distribution of retrogradely labeled cells following a bidirectional tracer injection in the ZI. a. injection site in the ZIr. b-f. labeled cells in frontal cortex, dots=charted labeled cells.

### ZI subcortical connections

#### Inputs

There were three injection sites, all of which were centered in the ZIr. The most prominent inputs were from the intralaminar nuclei, the medial hypothalamus, and the STN/VTA, the reticular formation (RF), and the PPN. Labeled cells in the intralaminar nuclei were concentrated in the dorsal centromedian nucleus (CMn) (Fig. 10a). Rostral to the ZI, labeled cells were located along the medial hypothalamus, primarily in the dorsomedial cell group, but extending into the ventromedial cell group (Fig. 10b). SN/VTA labeled cells were located primarily in the dorsal tier of the dopaminergic cell group (Fig. 10c). An anterograde injection into the dorsal SN/VTA resulted in labeled terminals in the ZI (Fig. 10ci) and is consistent with dopamine transporter-positive fibers coursing through and terminating in the ZIr [61]. Connections with the RF and PPN extended from the midbrain through the pons. In addition to these main inputs, clusters of labeled cells were located in the amygdala (amygdalo-hippocampal junction (Fig. 10d), in the central amygdala nucleus, and within the sublenticular extended amygdala), the dorsal raphe nucleus (DRN), pedunculopontine nucleus (PPN), and periaqueductal grey (PAG) (Fig. 10e). Finally, a few labeled cells were found in the medial dorsal nucleus of the thalamus and in the superior colliculus.

**Figure 10.**
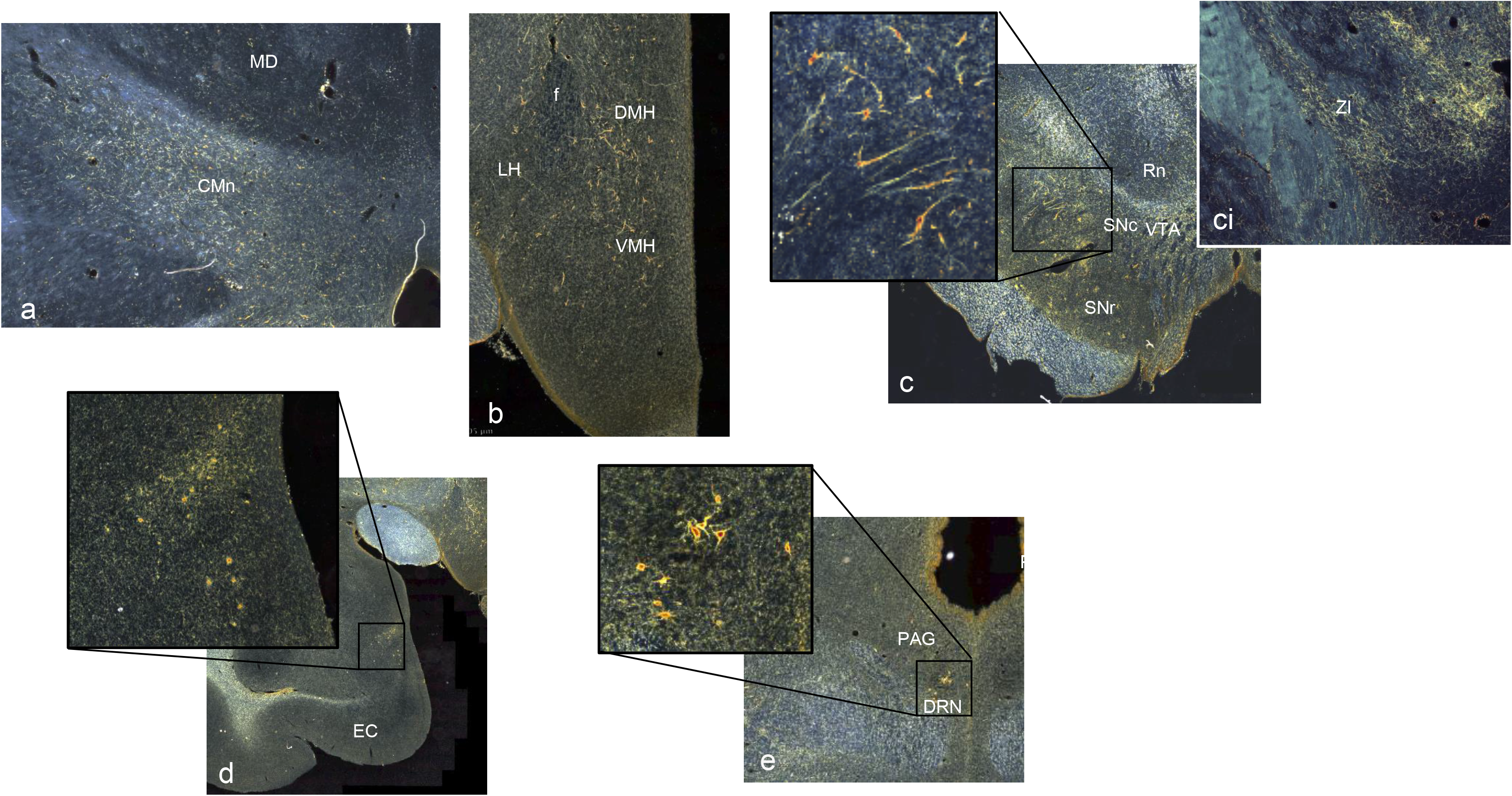
Photomicrographs illustrating labeled cells following a tracer injection into the ZIr (Fig. 8a). a. labeled cells and fibers in the intralaminar thalamic n. b. labeled cells in the medial hypothalamus. c. Labeled cells in the substantia nigra. ci=labeled of fibers in the ZIr following an anterograde tracer injection into the VTA. d. labeled cells in the amygdalo-hippocampal junction. e. labeled cells in the periaqueductal grey and dorsal raphe nucleus. CMn=centromedian nucleus, DMH=dorso-medial nucleus of the hypothalamus, DRN=dorsal raphe nucleus, EC=entorhinal cortex, LH=lateral hypothalamus, Md=medial dorsal nucleus of the thalamus, PAG= periaqueductal grey, Rn=red nucleus, SNc=substantia nigra, pars compacta, SNr=substantia nigra, pars reticulata, VMH=ventro-medial nucleus of the hypothalamus, VTA=ventral tegmental area, ZI=zona incerta.

#### ZIr outputs

The most prominent outputs from the ZIr are to the CMn, LHb, RF, and PPN. Additional projections are to the hypothalamus, SN/VTA, PAG, DRN, and SC. Fibers exit the ZI medially. Some axons travel dorsolaterally to enter the thalamus and travel caudally, terminating throughout the intralaminar nuclei, primarily in the CMn (see Fig. 10a). Axons continue caudally and dorsally, through the thalamus, and give rise to a large dense patch of terminals in the LHb (Fig. 11a). A retrograde injection placed into the LHb nucleus confirms this projection, with labeled cells throughout the ZIr (Fig. 11b). However, few labeled cells were found in the central ZI and none in ZIc. A second group of fibers exit the ZI and continue medially and caudally. Fibers either circumvent or pass through the red nucleus and form a dense bundle medial to the red nucleus which continues caudally. Dense terminals were located in the midbrain RF and PPN, with some fibers terminating in the PAG, and DRN. Finally, a third group of fibers turn ventrally from the ZI, arching around the medial STN, to enter and terminate in the dorsal SNc/VTA. Additionally, terminals were noted in the superior colliculus, but few or no axons were found in PFC/ACC cortex. Taken together, although the ZIr connections are widespread, the strongest inputs to the ZIr are from the PFC/ACC, CMn, medial hypothalamus, SN/VTA, RF, and PPN. The strongest outputs include reciprocal connections with the CMn, RF and PPN and an output to the LHb (Fig. 12).

**Figure 11.**
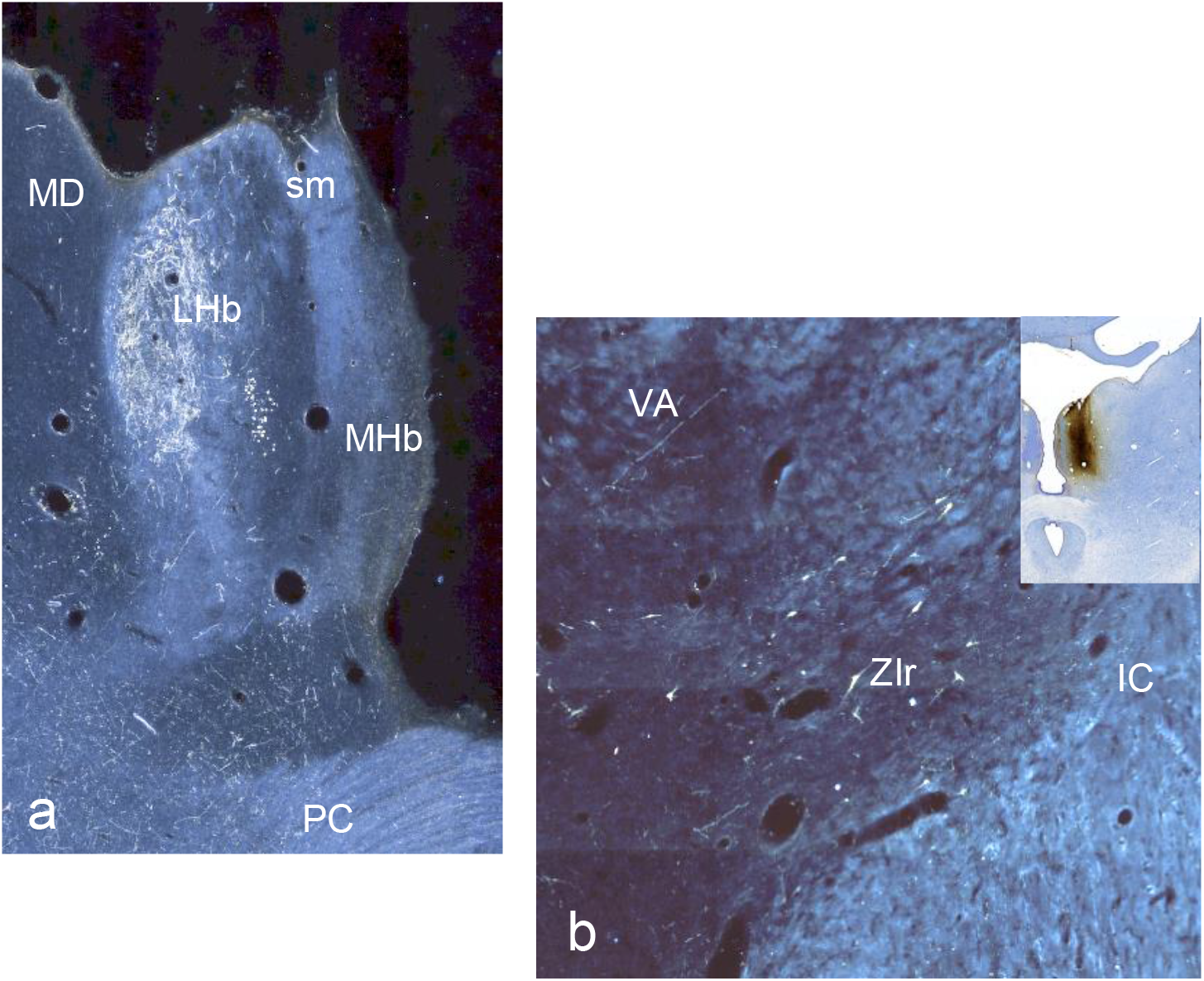
ZIr projections to the lateral habenula n. a. terminals following a tracer injection into the ZIr. b. labeled cells in the ZIr following a retrograde tracer in the lateral habenular n.(inset). IC=internal capsule, LHb=lateral habenula, MD=medial dorsal thalamic nucleus, MHb=medial habenula, LHb=lateral habenula, PC=posterior commissure, sm=stria medularis, VA=ventral anterior thalamic nucleus, ZIr=rostral zona incerta.

**Figure 12.**
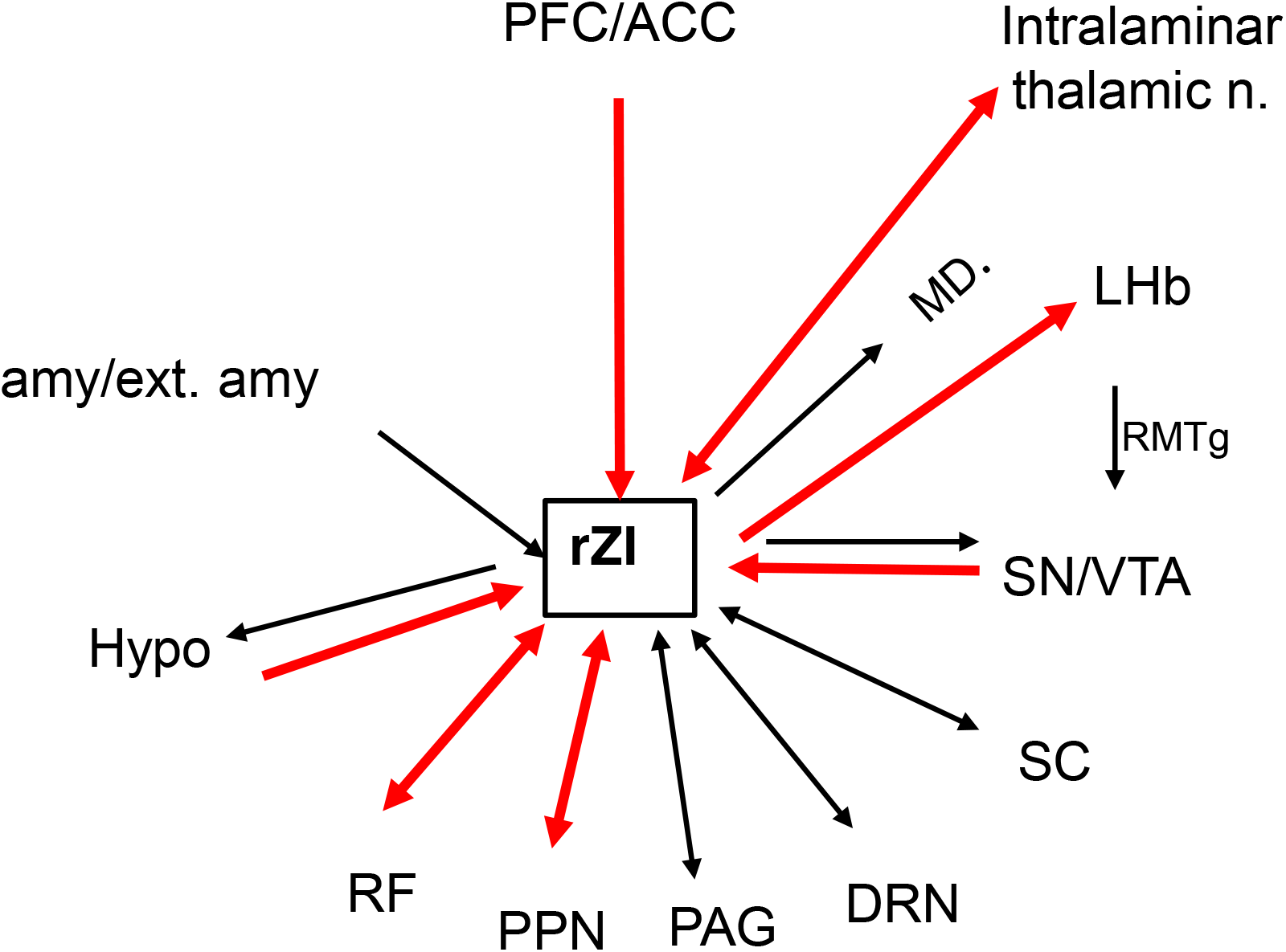
Schematic illustrating the main connections of the ZIr. Red arrows indicate the strongest connections.

## Discussion

The ZI is a highly conserved structure across species, including birds and reptiles [22, 52, 53, 85-89]. It is involved in several behaviors, including visceral function, arousal, attention, and locomotion. In mammals, the ZI is divided into four components, ZIr, ZId and ZIv, and ZIc, each with some distinct cytoarchitectural and chemoarchitectural features [22, 38, 51, 61, 90]. Rodent studies have focused primarily on the central and caudal portions of the nucleus, the ZIv/ZIc and ZId. These areas are linked to sensorimotor connections and function, and limbic connections and function, respectively. Few studies have directly examined connections of the ZI in nonhuman primates. The studies that have focused on ZI connections identified specific motor and sensory connections to the central or caudal ZI [53, 54, 55, 56, 89, 91–95]. Additional cortical inputs have been noted from the anterior and posterior cingulate cortices and from the anterior, ventromedial temporal cortex [56, 96–98]. Subcortical connections include: PAG, superior colliculus, interstitial nucleus of Cajal, pontine nuclei superior colliculus, interstitial nucleus of Cajal, pontine nuclei (PAG), and the spinal cord [99–103]. Finally, descriptions of dopaminergic terminals in the ZI [61] suggest, as in rodents, a dopaminergic input. Overall, these connections are consistent with those from homologous regions in the rodent. Here, for the first time, we identify the inputs to the ZI from higher cognitive areas in the primate PFC/ACC.

### Connections of the ZIr

The results demonstrate that the concentration of PFC/ACC terminals are located in the ZIr. At this level, there are few or no premotor and motor fibers. As the PFC/ACC fibers course through the ZI and terminate in the rostral part of the ZId, interfacing with axons from premotor areas. At the more caudal levels, as primary motor fibers enter the ZI, there are few or no PFC/ACC fibers. However, within the PFC/ACC projections, there appears to be little topography, with most fibers entering and overlapping in ZIr. The strongest PFC/ACC projections are derived from the dorsal and lateral PFC and the dorsal ACC, but not from the ventral and medial regions (OFC, subgenual ACC). The central ZI and ZIc receive the strongest inputs from premotor areas. Interestingly, while the ZI does receive input from motor control regions [53, 54, 55, 91–94], our results show that projections from primary motor cortex (M1) are remarkably sparse. Thus, there is a clear rostro-caudal organization of fronto-ZI projections, with PFC/ACC projecting rostrally, and motor control regions caudally. The main frontal inputs to the ZI arise from regions involved in higher cognitive and premotor regions with fewer inputs from limbic/affective-related regions and primary motor cortex. Other major inputs to the ZIr are from the CMn, medial hypothalamus, SN/VTA, RF, and PPN. Outputs are primarily reciprocal. Interestingly, one of the strongest ZIr projections is to the LHb, which is not reciprocal.

### Links to the basal ganglia

The dense output to the LHb along with connections with the SN/VTA link the ZI tightly with the BG, a system long associated with incentive-based learning. There are clear similarities in overall cortical connections of the ZIr with two key input BG structures, the striatum and STN: all three receive PFC/ACC projections. However, there is a critical difference. These projections to the striatum and STN are generally topographic. The VS and anterio-medial STN receive inputs from the OFC and ACC, the dlPFC and vlPFC project to the central striatum and STN, and motor control areas project to the posterior-lateral striatum and STN [66]. In contrast, all PFC/ACC projections to the ZI converge in the ZIr, with little topography. Thus, although there is some overlap between PFC/ACC inputs to the both the VS and STN, projections to the ZIr completely converge. Furthermore, as dlPFC, vlPFC, and dACC-striatal projections terminate in the caudate nucleus, they have little overlap with inputs from the amygdala, hypothalamus, PPN, and DRN, which are connected to the ventral striatum [104, 105]. Moreover, connections of the STN (in addition to cortex) are to basal ganglia structures, (primarily the SN/VTA and pallidum). Finally, neither the striatum nor the STN project directly to the LHb. Taken together with the other connections of the ZIr, these links into the basal ganglia system place the ZIr in a position to compute and modify incoming motivational and attentional signals, providing top-down inhibition (via GABA) to responses triggered by salient input.

### A ZIr connectional hub

The widespread PFC/ACC inputs to the ZIr coupled with its subcortical connections place the ZIr in a unique anatomic position to compile and modify information from a broad range of functions (Fig. 12). While the concept of hubs emerged in early discussions of brain networks [106], recent advances of network analysis have formalized the definition of hubs based on graph theory as a node with unusually high and diverse connections, which facilitates communication between functional modules [19, 20, 21]. Human resting-state functional magnetic resonance imaging (rsfMRI) studies have characterized large-scale distributed cortical networks and hubs within them [19, 20, 107–110]. Importantly, studies show strong links between the hard-wiring of a network, based on anatomic studies and connectivity identified with functional MRI [21]. However, few have focused on subcortical areas. Here we propose that the ZIr, based on its hard-wiring, may serve as a subcortical connectional hub that brings together elements of the BG, and other subcortical regions, with the PFC/ACC network hubs to mediate not only incentive-based learning but behavioral flexibility.

The ZIr shows two defining features of a hub: high degree-centrality (or connectivity) and high diversity. The high degree-centrality is reflected by the number of areas with dense projections to the ZIr. This includes several frontal cortical regions (specifically, dlPFC, vlPFC, rACC and dACC, and premotor cortex), diencephalic regions (intralaminar nuclei and hypothalamus), with additional inputs from the brainstem structures (SN/VTA, PAG, PPN, DRN, SC). Importantly for a hub definition, these inputs also represent a highly diverse set of connections. The strongest cortical inputs to the ZIr mediate a broad set of behaviors that are central for cognitive control and behavioral flexibility. The rACC sits at the connectional intersection of the motivation and action control networks, an important position in the transition from valuation through choice to action, particularly in situations of uncertainty [81, 111–115]. It is also considered a hub and one of the main anchors within the default mode network [18, 116, 117]. The dACC (caudal to the genu) is more tightly connected with the action network consisting of motor control areas, including frontal eye fields and premotor areas [118, 119], and is associated with motor planning and action execution [120, 121]. The dlPFC is particularly linked with action associated with working memory tasks [122–124]. The vlPFC has been implicated in memory retrieval, reversal learning, and overall behavioral flexibility, [7, 8, 125–129]. Importantly, different vlPFC areas are linked to the three attention networks, the ventral and dorsal attention networks and the salience networks. It can also be considered a hub, important for switching between the three attentional networks [130]. The ZIr thus receives inputs from cortical hubs, fulfilling the key criteria for a connectional hub. Diencephalic and brainstem connections provide bottom-up information about the internal and external world.

#### The role of the ZI in modulating behavior

The ZI is involved in the modulation of fear responses, extinction, and defense mechanisms [28–30], reinforcing motivational drive [33, 34], and novelty seeking [27, 56], all of which require quick behavioral adaptation and decision-making skills in response to salient stimuli. It is an important node in modulating behaviors, particularly those for survival [26]. Most behavioral studies manipulate the central or caudal ZI, the sensory integration and motor control regions [27, 29, 30, 33, 35, 56], However, in NHPs, the ZIr is more likely involved in the higher decision-making aspects of these behaviors. Indeed, a few rodent studies have shown the importance of the ZIr in modulation of appetitive, defense and predatory behaviors [28, 34, 35]. The relatively dense projections from the dlPFC, vlPFC, rACC, and dACC, suggest that the ZIr in primates has a cognitive role in top-down behavioral modulation. These inputs, while concentrated in the ZIr extend into the rostral central region, thus in a position to interact with inputs from premotor areas. The dense direct projection to the LHb, combined with connections with the SN/VTA, is particularly interesting as it places the ZIr in an optimal position to uniquely contribute to modulation of value-based predictions [59, 131, 132]. Coupled with other ZIr subcortical connections, including the amygdala, hypothalamus, PAG, and DRN, the ZIr is in a central position to integrate cortical and subcortical information, functioning as a connectional hub in a network involved not only in value encoding and reinforcement learning, but also behavioral flexibility.

#### Implications for DBS

DBS is an effective treatment for movement disorders, including Parkinson’s disease (PD), Tourette syndrome, dystonia, and tremor. Stimulation primarily targets myelinated axons with the aim of interrupting abnormal information flow within specific networks [133, 134, 135, 136, 137, 138, 139]. Electrode placements for movement disorders target the basal ganglia motor circuit, and most commonly placed dorsolaterally in the STN. More recently, several studies have shown that the ZI is also an effective DBS target for motor disorders. Specifically, the target is the ZIc, dorsal to the motor sector of the STN and ventral to the motor thalamus. Indeed, it has been suggested that this target may be superior to the STN for movement disorders [140–146].

DBS is also a promising therapeutic approach for several treatment-resistant psychiatric diseases, including OCD [10: Figee, 2013 #14466, 11-16, 63, 147, 148]. The electrode targets for OCD are more wide-spread than those for PD, but nonetheless focus on the cortico-basal ganglia network, specifically the OFC/ACC-BG pathways. The main targets are the ventral ALIC, the VS, and the ventromedial STN (limbic region). Additional targets that have achieved some successes are the VTA, inferior thalamic peduncle (ITP) and the bed nucleus of the stria terminalis [149–152]. Despite efforts to refine the electrode targets for OCD, clinical outcomes following treatment have not improved significantly and remain at approximately 50%.

Patients with OCD are impaired in guiding behavior to optimize the balance between positive and negative outcomes, reflecting impairments in behavioral flexibility and decision-making [153]. Brain regions involved in behavioral flexibility are not limited to only the OFC and ventromedial regions, but also include more dorsal and lateral areas, i.e., the dorsal ACC and vlPFC [128] [129]. Additionally, the dlPFC is central for cognitive control. Terminals from these three cortical regions converge within the ZIr. This convergence is well positioned to modify responses from bottom-up areas important for rapid survival responses to provide a quick pause to better assess the likely outcome. These connections coupled with portals into the basal ganglia system provide a mechanism to impact directly on action outcomes to salient stimuli. This places the ZIr hub at the crossroads for modulating behavioral flexibility.

Here, we propose the ZIr as a potentially effective DBS target for OCD (Fig. 13). An electrode placed in the ZIr would modulate most of the connections captured at the other sites [12] [14], but with some important differences. The goal of the ALIC, VS, STN, VTA, ITP targets is to capture primarily the ventral cortico-basal ganglia system, the ‘limbic/affective’ circuit, but not the cognitive control circuits. That is, the OFC/ventral ACC fibers and their basal ganglia connections (VS, thalamus, STN, VTA, etc.), but not the dorsal ACC (dACC), vlPFC, medial PFC (mPFC), and dlPFC. As described above, PFC/ACC fibers and terminals in the ALIC, VS or STN, are topologically organized [66, 82, 154]. The ALIC, VS, STN, VTA DBS electrodes for OCD are placed to capture the ventral PFC/ACC fibers: in the ventral capsule, VS, anteromedial STN, and VTA [14]. The ITP carries ventral frontal cortical fibers to the thalamus and is targeted to involve OFC connections. The BNST target is placed adjacent to the IC at the junction with the caudate nucleus. This position is similar to that in which PFC/ACC fibers change orientation within the IC and begin to exit to the thalamus and likely captures many of the PFC/ACC fibers as they exit to the thalamus.

**Figure 13.**
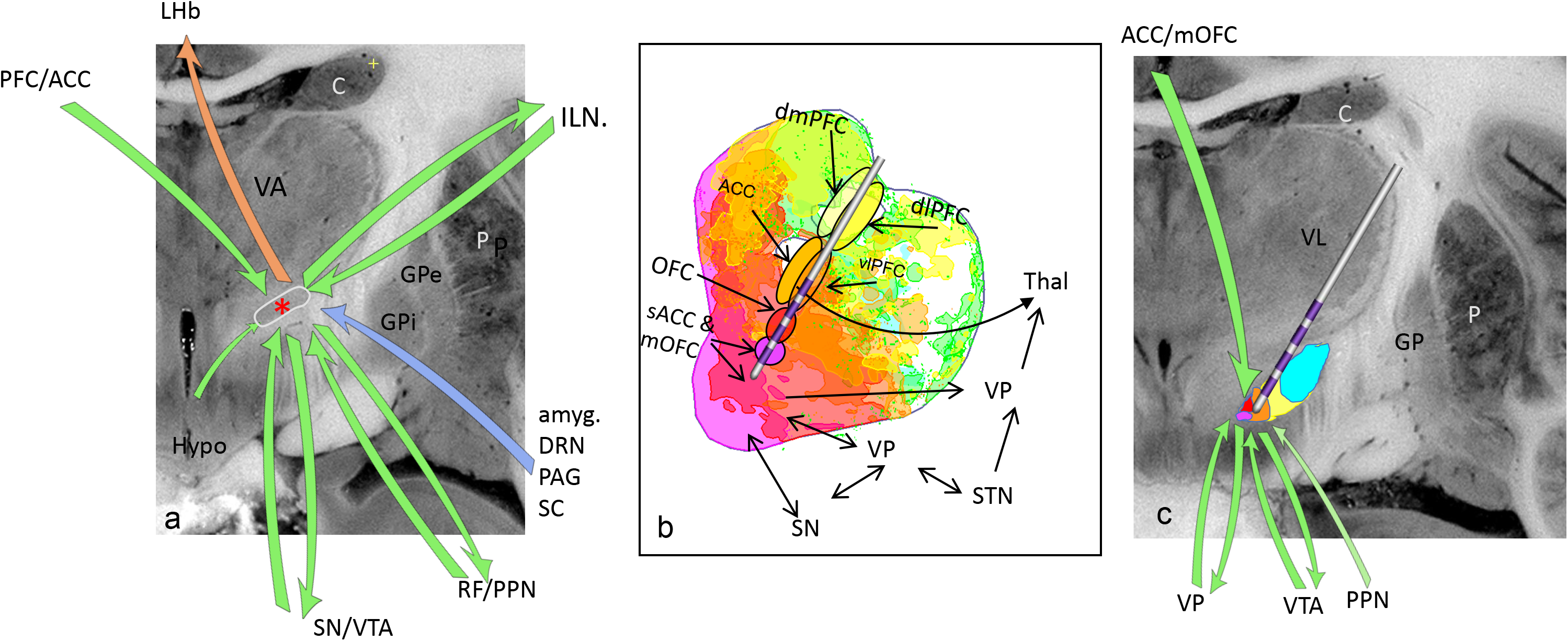
Potential ZIr DBS site and its connections compared to the most common sites for OCD (VS/ALIC & STN). a. ZIr site (red asterisk) and the major (green) other (blue) connections, and projection to the LHb (orange). Note: Connections of the ZIr include convergent inputs from all PFC/ACC, compared of topography cortical fibers through the ALIC and inputs to the VS &STN. ZIr connects with both BG structures and non-BG structures. Red asterisk=Potential DBS site. b. Combined ALIC and VS sites. The electrode typically targets the ventral capsule/VS area. Both sites primarily involve ACC/OFC fibers, and/or ventral basal ganglia structures. c. STN site targets primarily ACC/OFC fibers and ventral basal ganglia structures. amyg.=amygdala, C=caudate nucleus, dlPFC=dorsolateral PFC, dmPFC=dorsomedial PFC, DRN=dorsal raphe nucleus, GP=globus pallidus, ILM=intralaminar nuclei, LHb=lateral habenula, mOFC=medial OFC, PAG= periaqueductal grey, PFC/ACC=prefrontal cortex/anterior cingulate cortex, OFC=orbitofrontal cortex, P=putamen, PPN=pedunculopontine nucleus, RF=reticular formation, sACC=subgenual ACC, SC=superior colliculus, SN=substantia nigra, STN=subthalamic nucleus, thal=thalamus, VL=ventral lateral nucleus, vlPFC=ventrolateral PFC, VP=ventral pallidum, VTA=ventral tegmental area,

An electrode placed at the ZIr would capture more specifically the dorsal and lateral PFC axons, although some ventral connections would also be included. This placement would compare to a more dorsal ALIC, striatal or central STN electrode placement. However, these positions then would not include other potentially important bottom-up connections, including the amygdala, hypoth, PPN, PAG, DRN, etc. These structures are primarily involved in the ventral cortico-basal ganglia system, but not the dorsal system. Taken together, a ZIr target provides a unique opportunity to link cognitive control regions that are in a position to more directly modulate bottom-up inputs. Importantly, the ZIr target would also directly impact on the LHb, a DBS site proposed for depression [155], and the serotonin ascending system (pharmacological therapeutic target for OCD).

A key requirement for targeting this system in therapeutic interventions is the ability to localize it accurately in the human brain. New methods for localizing subcortical structures are effective for identifying small nuclei, including the ZI [156]. Here we have shown the feasibility of reconstructing the projections of the ZI using tractography on human diffusion MRI data acquired in vivo at sub-mm resolution. These results show remarkable agreement with the NHP tracing, but require a very long, multi-session diffusion MRI protocol that is impractical for routine use in patients. One possibility for reducing the acquisition time is to increase the signal-to-noise ratio at sub-mm resolution, e.g., by going to higher field strengths (7T or higher). Furthermore, we have previously shown that, if tracts can be annotated manually on high-quality diffusion MRI data, they can be used to train an automated tractography algorithm to reconstruct the same tracts in lower-quality data [157]. While imaging the projections of the ZI in individual patients will require further development of data acquisition and analysis techniques, prior work has shown that high-quality normative data may be better suited than low-quality individual data for DBS targeting [158]. Thus, we are making available the pathways that were annotated in a sub-mm human dataset for the present work, to facilitate further research on targeting the ZI for neuromodulation.

## Acknowledgments

This work was supported by the National Institute of Health (Grants Nos. MH106435 and MH045573 to Dr. Haber).

